# Five years investigation of female and male genotypes in Périgord black truffle (*Tuber melanosporum* Vittad.) revealed contrasted reproduction strategies

**DOI:** 10.1101/073650

**Authors:** Herminia De la Varga, Françis Le Tacon, Mélanie Lagoguet, Flora Todesco, Torda Varga, Igor Miquel, Dominique Barry-Etienne, Christophe Robin, Fabien Halkett, Francis Martin, Claude Murat

## Abstract

The Périgord black truffle (*Tuber melanosporum* Vittad.) is a heterothallic ascomycete that establishes ectomycorrhizal symbiosis with trees and shrubs. Small-scale genetic structures of female genotypes in truffle orchards are known, but it has not yet been studied in male genotypes. In this study, our aim was to characterize the small-scale genetic structure of both male and female genotypes over five years in an orchard to better understand the *T. melanosporum* sexual reproduction strategy, male genotype dynamics, and origins. Two-hundred forty-one ascocarps, 475 ectomycorrhizas, and 20 soil cores were harvested and genotyped using microsatellites and mating type genes. Isolation by distance analysis revealed pronounced small-scale genetic structures for both female and male genotypes. The genotypic diversity was higher for male than female genotypes with numerous small size genotypes suggesting an important turnover due to ascospore recruitment. Larger and perennial female and male genotypes were also detected. Only three genotypes (1.5 %) were found as both female and male genotypes (hermaphrodites) while most were detected only as female or male genotype (dioecy). Our results suggest that germinating ascospores act as male genotypes, but we also proposed that soil mycelium could be a reservoir of male genotypes.

## Introduction

The Périgord black truffle, *Tuber melanosporum,* is an ascomycete species that forms ectomycorrhizal symbiosis with trees and shrubs. For forty years, large-scale inoculation of tree seedlings with *T. melanosporum* ascospores has been used in nurseries to produce in France >300,000 symbiotic plantlets yearly (Chevalier and Grente, 1978; Murat, 2015). As a consequence, truffle orchards are now found worldwide (Europe, Australia, USA, South America, South Africa, and New Zealand). In these truffle orchards, the production of ascocarps (fruiting bodies issued from sexual reproduction) faces several problems, one of them being the initiation of sexual reproduction (Chen et al., 2016). The factors involved in truffle sexual reproduction are difficult to investigate due to the impossibility of manipulating truffle *in vitro* (Le Tacon et al., 2015). Because this information is still missing, clear guidelines for truffle orchard management are still missing.

*Tuber melanosporum* is a heterothallic species having two mating type idiomophs (*MAT1-1* and *MAT1-2*) (Martin et al., 2010; Rubini et al., 2011b). Mating in heterothallic ascomycetes results from the union of female gametes (ascogonia) and male gametes of opposite mating types. Male gametes can be formed in antheridia (haploid structure containing male gametes). Conidia (haploid asexual spores) or haploid hyphae can also act as male gametes (Glass and Kuldau, 1992; Leslie and Klein, 1996). In heterothallic species, male and female gametes can be formed by the same haploid mycelium (one genotype) defining the so-called hermaphroditism or monoecy. In another situation, male and female gametes are formed by distinct male and female haploid mycelium (two different genotypes) corresponding to dioecy. In *T. melanosporum,* the female gametes are ascogonia produced by the haploid mycelium forming the ectomycorrhizal root tips. The ascogonia produced from this haploid mycelium can be either *MAT1-1* or *MAT1-2* (Rubini et al., 2011b; Murat et al., 2013; Taschen et al., 2016). They give birth to the main structure of the ascocarp (the peridium and the unfertile tissues of the gleba) (Fig. 1). The origin of male gametes is still unknown, but it has been suggested to be germinating sexual ascospores, conidia, or persisting soil mycelium (Fig. 1) (Le Tacon et al., 2015; Taschen et al., 2016).

**Fig. 1.**
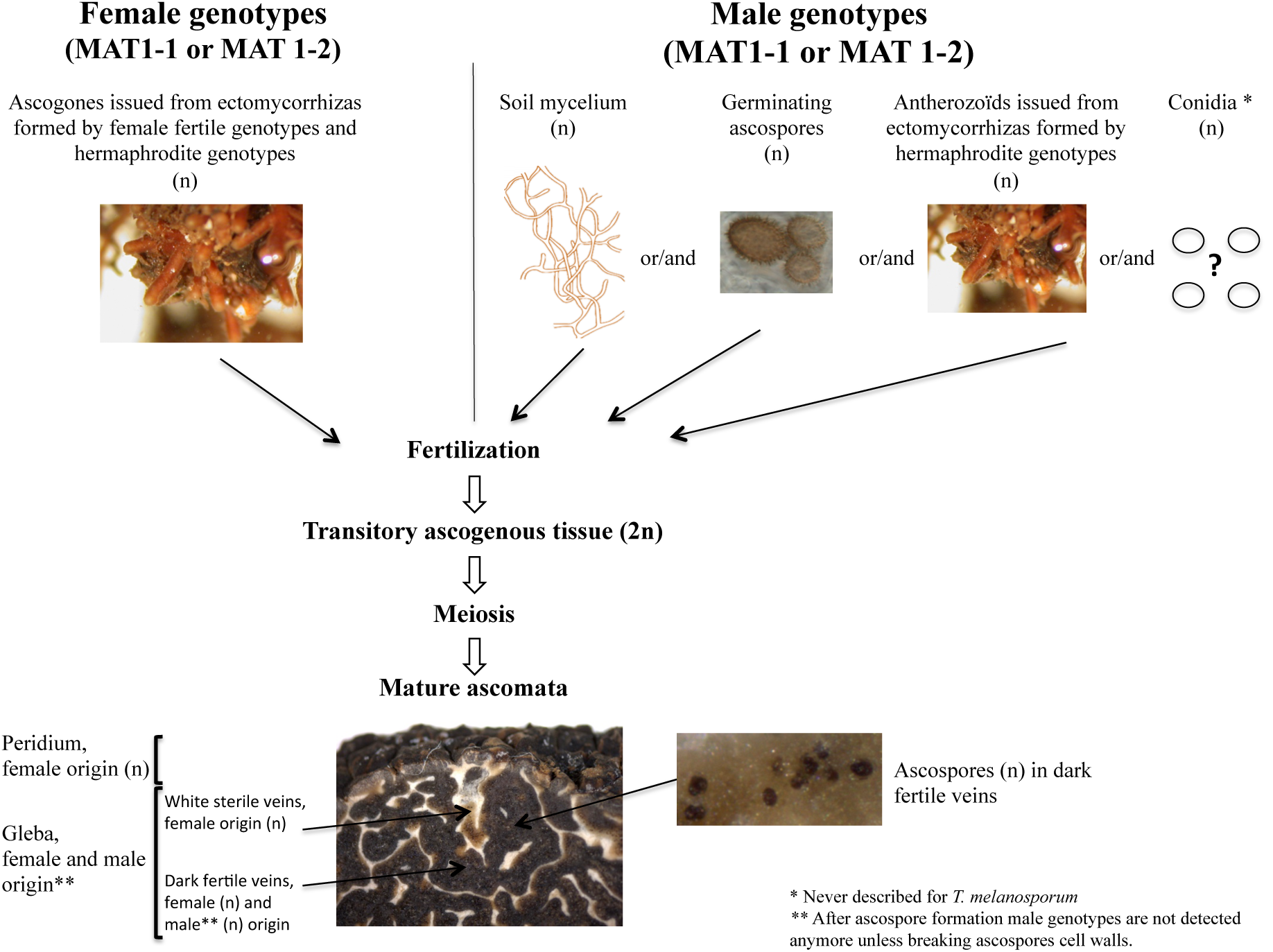
Schematic representation of sexual reproduction for the heterothallic ascomycete *T. melanosporum.* In order that sexual reproduction occurs female and male genotypes needs to mate. In nature, the female genotype is found in the host tree root system as ectomycorrhiza, although the male genotype could have different origins: germinating ascospores, soil mycelium (free-living or associated to ectomycorrhiza for hermaphrodite genotype), or conidia. Conidia have never been observed for *T. melanosporum,* but they have been described in other *Tuber* species. After the fertilization, a diploid transitory phase occurs (that cannot be detected in mature ascocarps), followed by a meiosis phase that will end in the formation of a mature ascocarp. The structures of ascocarps (peridium, gleba. and ascospores) are represented in the photographs.

One approach to help understand the truffle's sexual reproduction is to characterize small-scale genetic structures for both female and male genotypes. Indeed, small-scale genetic structure depends on the distribution of genotypes in population and results from a combination of their propagation modes and demographic processes. This can be analysed by characterizing the persistence and size of the genotypes (Douhan et al., 2011). The isolation by distance (IBD) theory predicts that a small-scale genetic structure accounts for gene dispersion capacity (Vekemans and Hardy, 2004). The small-scale genetic structure of female *T. melanosporum* genotypes was studied by genotyping with microsatellites and mating type genes of ectomycorrhizas and ascocarps (Rubini et al., 2011a; Murat et al., 2013; Taschen et al., 2016). In natural stands and truffle orchards, a pronounced genetic structure was observed. Moreover, ectomycorrhizas and female genotypes formed patches containing only one of the two mating types (spatial segregation). This particular spatial distribution raises the questions about the origins and dynamics of the male gametes since a high density of ascocarps was not observed at the boundary of patches with opposite mating types. This observation also questioned the existence of hermaphroditism (capacity of one genotype to form either male or female gamete) for *T. melanosporum* (Fig. 2). Inside these mono mating type patches, up to 10 female genotypes were detected; some were perennial (found during several seasons and forming genotypes with size up to 4.7 m), while others were detected only in one season and formed genotypes less than one meter (Murat et al., 2013; Taschen et al., 2016). This pattern suggests a rapid turnover of the non-perennial female genotypes. Recently, Taschen and colleagues (2016) also surveyed male genotypes in five brulés (areas without vegetation surrounding trees that are substantially mycorrhized by *T. melanosporum*) located on three plantations and two natural sites. The genetic diversity was higher for male than female genotypes, and most of the male genotypes were detected once only. Only a single male genotype occurring more than one year was identified. However, the male genotype small-scale genetic structure could not be assessed in that study.

**Fig. 2.**
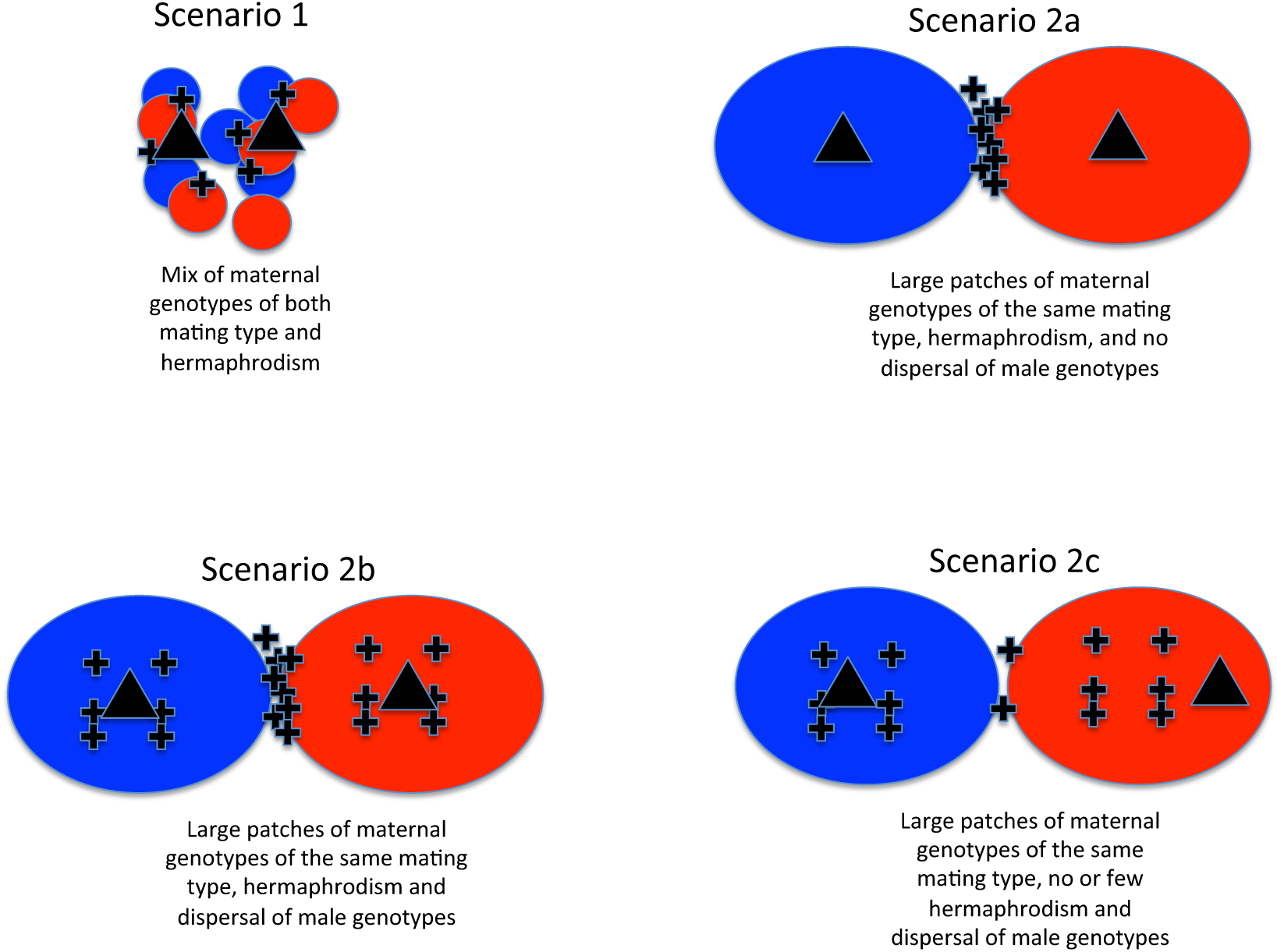
Theorical scenarios for the localisation of ascocarps for an ectomycorrhizal heterothallic ascomycete according to the sexual strategy. The trees are indicated as triangle and ascocarps as crosses. Scenario 1 considered that on the root system female genotypes of both mating type are intermixed and they are hermaphrodites. This scenario is not true for *T. melanosporum* since genotypes of the same mating type tend to aggregate (Rubini et al., 2011a; Murat et al., 2013; Taschen et al., 2016 and this study). Scenarios 2a-c considered that the aggregation of genotypes in the root system was observed in *T. melanosporum* and less or more hermaphroditism and male dispersion.

The objective of the present study was to characterize the small-scale genetic structure of both male and female genotypes in order to clarify the *T. melanosporum* sexual reproduction strategy and, more particularly, the origin and the behavior of the male genotypes. This work addresses several main points: 1) the small-scale genetic structure of the male genotypes and their evolution according to the time (perennial versus transitory male genotypes); 2) the localisation in the field of male genotypes in ascocarps, ectomycorrhiza, and soil; 3) the possible origin of the male genotypes; and 4) the sexual reproduction strategy (hermaphroditism versus dioecy). To achieve these goals, we used a 5-year sampling of ascocarps, ectomycorrhizas, and soils under seven productive and contiguous trees in a 25-year-old established truffle orchard in northeastern France. We determined the genotypes of both female and male ascocarp partners in addition to that of mycorrhizas and the mating type present in the soil samples.

## Results

### Ascocarps and ectomycorrhiza samplings

A total of 241 ascocarps were harvested during five consecutive truffle production seasons (from November to March of the following year) from 2010/2011 to 2014/2015 (Table 1; Fig. S1). The ascocarps were randomly distributed around the productive trees and could be also found at four to five meters from the trunk in and between the zone of root extension (Fig. S1). The productive area under each tree extended slowly over five years.

**Table 1.**
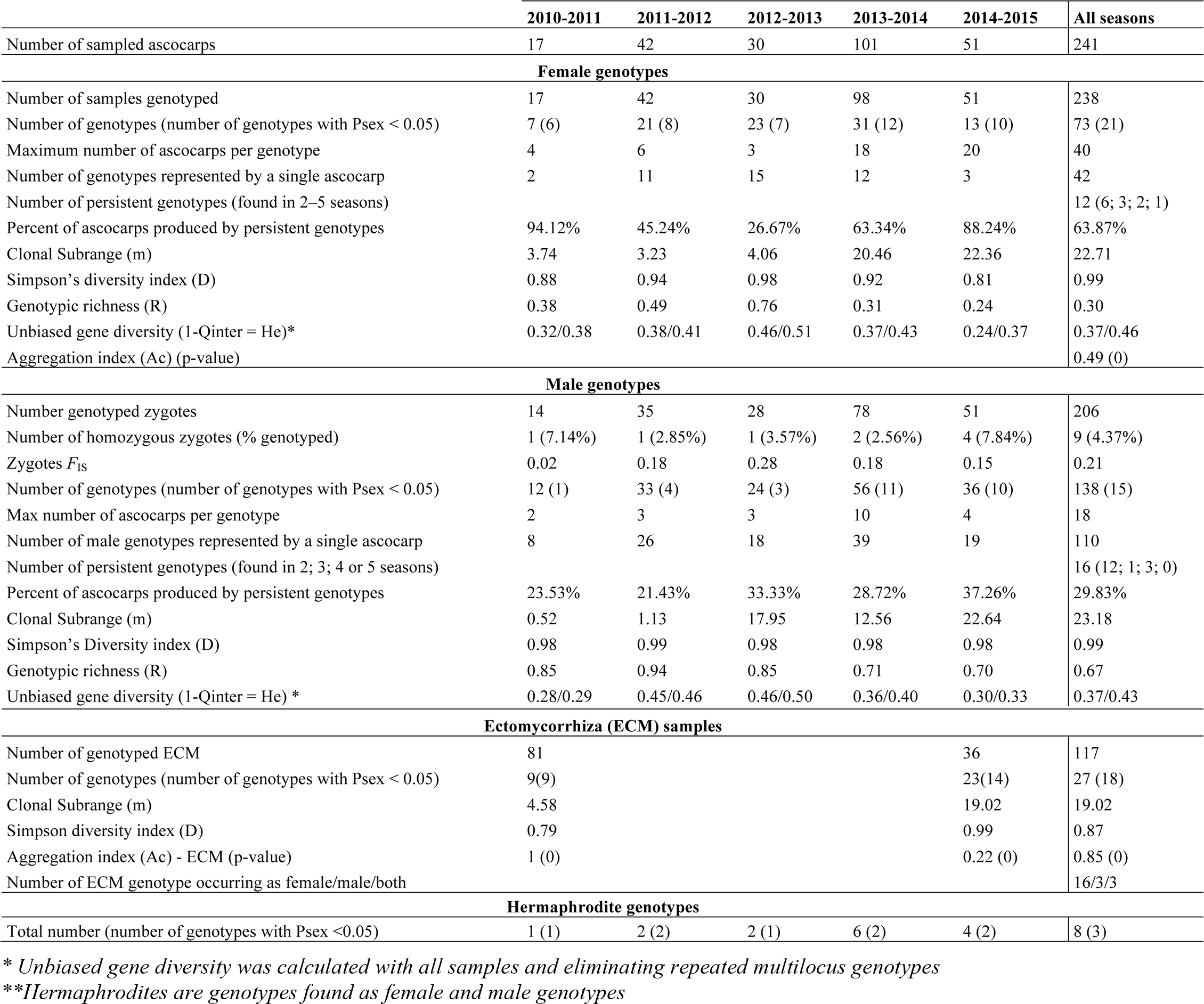
Sampling details, genotypic, and genetic diversity for female and male genotypes and ectomycorrhiza over five years from the Rollainville orchard.

Four-hundred seventy-five ectomycorrhizal root tips were collected (205 in 41 root samples with five *T. melanosporum* morphotypes each in 2011 and 270 in 45 root samples with six *T. melanosporum* morphotypes each in November 2014) (Fig. S1). Only 37 ectomycorrhiza samples did not amplify with the *T. melanosporum* specific primers, and the presence of *T. melanosporum* ectomycorrhizas was not confirmed on five root samples (two in 2011 and three in 2014). For microsatellite genotyping, only one *T. melanosporum* ectomycorrhizal root tip per root sample was selected.

### Characterization of female genotypes (gleba and ectomycorrhiza)

The female ascocarp genotypes (gleba) were successfully obtained for 238 out of the 241 ascocarps (Table 1 and Table S1). Seventy-three female multilocus genotypes (called hereafter genotypes) with significant Psex (probability that two identical genotypes originated from the same genet) value were found during the five years (Table 1, Tables S1 and Table S2). Twelve genotypes were present in more than one season (Table S1). These genotypes accounted for 26% to 94% of the ascocarps for seasons 3 and 1, respectively, for a total of 154 truffles (64% of the total harvested ascocarps over the five seasons). One persistent genotype (R002) was found throughout all of the seasons under the same tree (F11, area 3), representing 15% of the total ascocarps harvested (Fig. S2 and Table S1). Another genotype (R021) fructified under the A11 tree (in area 1) for three seasons and represented 16 % of the total harvested ascocarps (Fig. 3 and Fig. S1). The maximum size (clonal subrange) of the female genotypes was 22.36 m, which corresponded to the R002 genotype found in areas 1 and 3 of the truffle orchard (Table 1 and Fig. S2). Most female genotypes (58%) were found in only one ascocarp (Table S1).

**Fig. 3.**
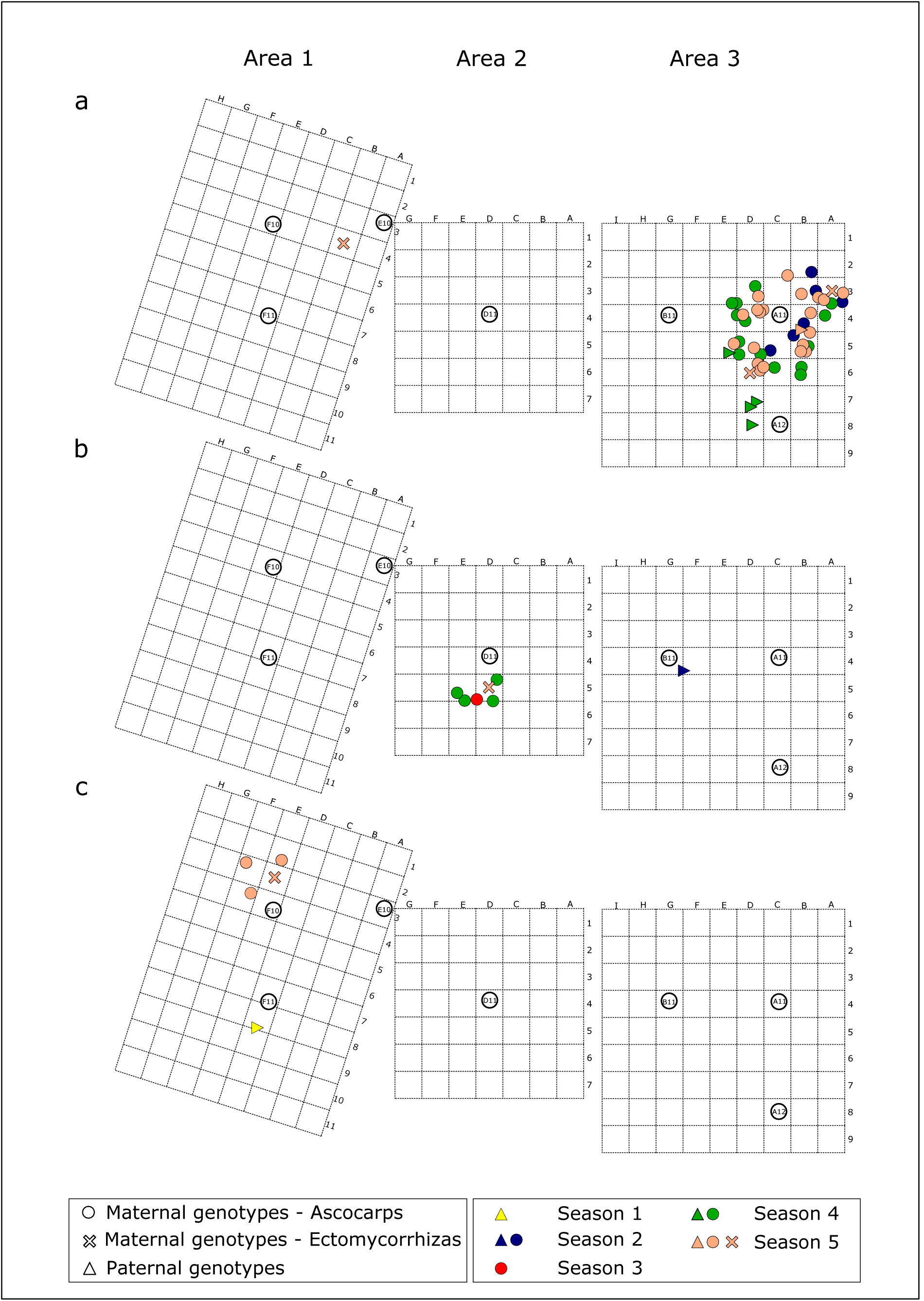
Map of distribution of the hermaphrodite genotypes R021, R060, and R068 from the Rollainville orchard. Representation of the distribution of the hermaphrodite genotypes along the different seasons from the 2010-2011 (season 1) to the 2014-2015 (season 5) and in the different areas of the orchard. a. Distribution of genotype R021. b. Distribution of genotype R060. c. Distribution of genotype R068. Dots represent female ascocarp genotypes, crosses represent ectomycorrhiza genotypes and triangles represent paternal genotypes. Each season is represented by a different colour.

One-hundred seventeen ectomycorrhizas representing nine and 14 genotypes for 2010 and 2014 samplings, respectively, were genotyped (Table 1 and Table S1). Five genotypes (R001∙– R004 and R007) were found in both ectomycorrhizal samplings in 2010 and 2014 (Table S1). The maximum ectomycorrhiza genotype size was 19.02 m for R021 (Table 1 and Fig. 3). Eighty-six percent of ectomycorrhiza genotypes were also detected as female genotypes in ascocarps, and were represented in 16 genotypes with significant Psex values (Table S1). Similarly, 86% and 70% of the female genotypes were also detected in ECM genotypes for seasons 1 and 5, respectively.

When considering only the mating type locus, the aggregation index (Ac) was 0.49 (p value = 0) for female genotypes, indicating that genotypes of the same mating type tend to aggregate (Table 1). In the orchard, large patches from 5 to 20 m^2^ of different genotypes of the same mating type for the root colonization were observed (Fig. S3). A single tree can be colonized by one patch (such as E10 and B11) or by two contiguous patches of opposite mating types (F10, F11, D11, A11, and A12). Tree D11 harboured maternal genotypes and ectomycorrhizas of *MAT1-2* with the exception of one ectomycorrhiza sampled in square F5 formed by *MAT1-1* mycelium (Fig. S3 and Table S1). This genotype was not detected in male genotypes (see below). At the contact zone between patches of opposite mating types, no differences in the ascocarp density were observed when compared to density within the patch (Fig. S1 and Fig. S3).

### Characterization of male genotypes

We successfully genotyped 206 zygotes (a mix of female and male genotypes in ascocarps coming from the mate of female and male gametes) out of the 241 ascocarps (86.5 %; Table S1). By subtracting the female genotype, we were able to reconstruct male genotypes. A total of 138 male genotypes were found (Table 1 and Table S1). Nine zygotes (4%) were homozygous for all microsatellite loci (only mating type locus is heterozygous). In zygotes, a significant heterozygote deficit was observed with *F*_IS_ values of 0.02 and 0.28 for seasons 1 and 3, respectively (Table 1). The relationship between female and male genotypes in zygotes was investigated using the kinship coefficient calculation. The kinship coefficient varied from −0.5 to 1.5 with a mean value of 0.25, indicating most of the female and male genotypes in zygotes were genetically close (Fig. S4). Indeed, a kinship coefficient of 0.25 might correspond to full-sibs (Loiselle et al., 1995). However, the kinship coefficient was negative for 26% of the zygotes, indicating that in those cases female and male genotypes were not genetically related.

Most of the male genotypes (75%) were transitory (found only in one ascocarp) (Table S1). Only eight persistent male genotypes with significant Psex values were found. These persistent genotypes produced 21% to 37% of the ascocarps (Table 1). Sixteen male genotypes were found during several seasons and five (only one genotype with significant Psex) were found under different trees (Table S1). One genotype (R102) was detected for four seasons under A11 where it fertilized 8.7% of the ascocarps (Fig. S5). The maximum male genotype size was 22.64 m for genotype R102 with detection of most of them in one single ascocarp (Table 1).

### Small-scale genetic structure for both female and male genotypes

The genotypic richness was always higher for male genotypes (from 0.70 to 0.85) than for female genotypes (from 0.24 to 0.76). Depending on the season, the Simpson’s diversity indices ranged from 0.81 to 0.98, 0.98 to 0.99, and 0.79 to 0.99 for female, male, and ectomycorrhizal genotypes, respectively (Table 1). The inter-individual diversity (1-Qinter), also called unbiased gene diversity (He), ranged from 0.24 to 0.46 and 0.28 to 0.46 for female and male genotypes, respectively (Table 1). The small-scale genetic structure of the *T. melanosporum* population was assessed using an IBD analysis with GenePop software. In IBD, the slope indicates dissemination capacities (the higher the slope, the more the dissemination capacities are reduced). Using the culled dataset, a significant genetic structure was detected for both female and male genotype; the slope was higher for female than male genotypes (0.045 versus 0.039), but not significantly different (Fig. 4).

**Fig. 4.**
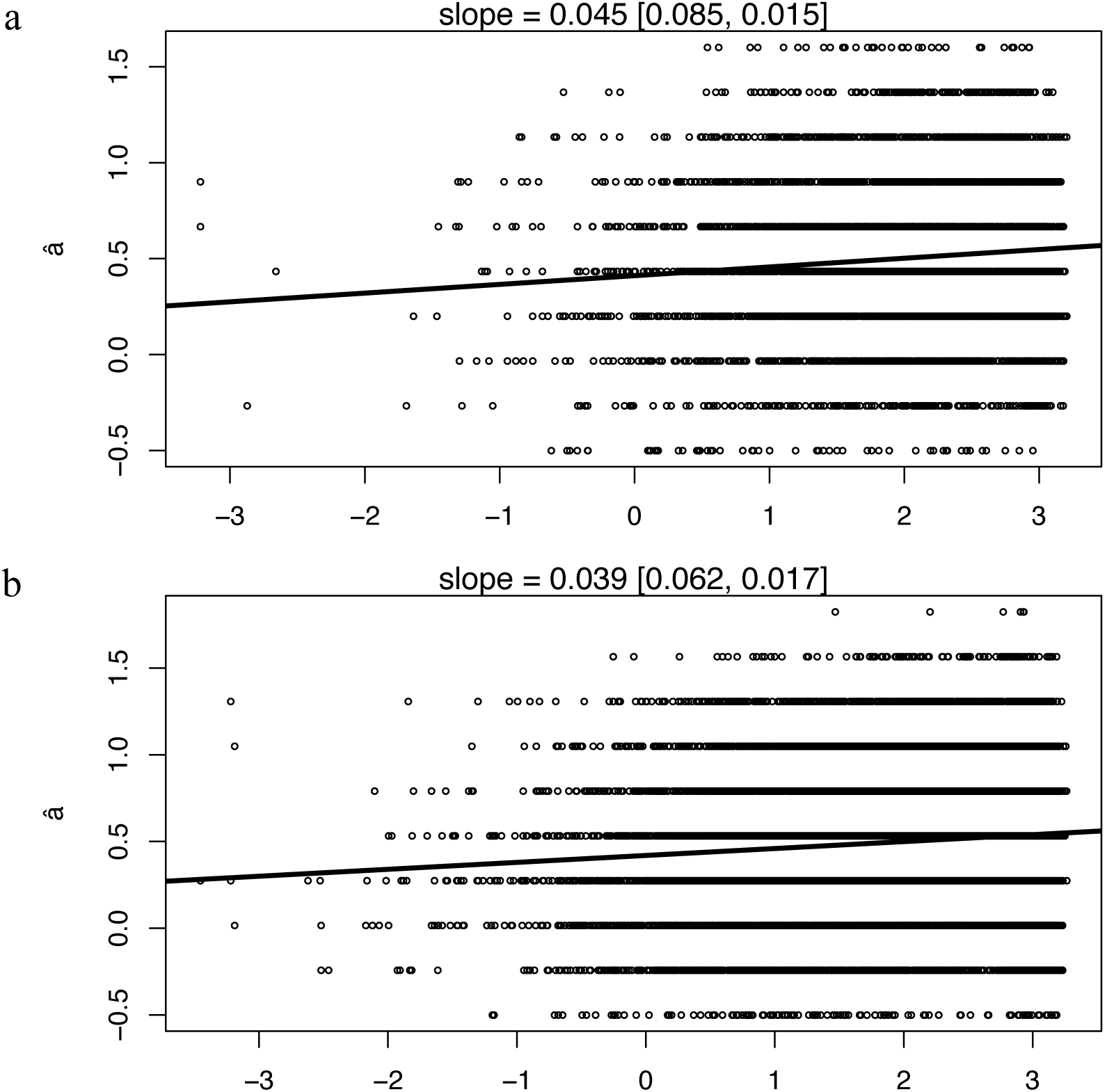
Isolation by distance analysis (IBD) representing genetic *versus* geographic distance for female and male genotypes. IBD analyses obtained with female (a) and male (b) genotypes using culled data to avoid oversampling of the same genotype. The slope with 95% interval confidence is indicated for each graph.

### Quantity of mating type myceliums in soil samples

In the 20 soil cores harvested in May 2015 (Fig. S1), both mating types were detected as mycelium in 16 soil samples (Table 2). In one soil sample, neither of the both mating types was detected, while in two soil samples only *MAT1-1* was found, and in one soil sample only, *MAT1-2* was detected. The quantity of mycelium ranged from 0 to 7.4 mg of mycelium per g of soil and 0 to 40.49 mg of mycelium per g] of soil for *MAT1-1* and *MAT1-2,* respectively. With two exceptions (squares E4 and E8 in area 1), the most frequent mycelium corresponded to that of female and ectomycorrhiza genotypes.

**Table 2.** Quantity of soil mycelium of both mating types in mg of mycelium/g of soil. The position in the truffle orchard is indicated in addition to the maternal mating type in the sampling square. For squares without ascocarps, the most frequent mating type of maternal tissue in the surrounding square is indicated (identified by an asterisk).

| Area | Square | Female MAT | *MAT 1-1* | *MAT 1-2* |
| --- | --- | --- | --- | --- |
| 1 | A4 | 2 | 0.428 | 16.37 |
| 1 | A8 | 2* | 0.402 | 21.58 |
| 1 | C3 | 2* | 0.444 | 18.22 |
| 1 | C6 | 2 | 0.408 | 23.03 |
| 1 | E4 | 1 | 0.753 | 40.49 |
| 1 | E8 | 1* | 0 | 0.021 |
| 1 | G3 | 1 | 6.14 | 0 |
| 1 | G6 | 1* | 7.4 | 0.04 |
| 2 | B4 | 2 | 0.535 | 25.39 |
| 2 | D2 | 2 | 0 | 0 |
| 2 | D6 | 2* | 0.11 | 5.6 |
| 2 | F4 | 2* | 0.022 | 1.13 |
| 3 | A2 | 2 | 0.171 | 9.51 |
| 3 | B7 | 2* | 0.179 | 9.009 |
| 3 | C4 | 2 | 0.567 | 31.56 |
| 3 | D5/6 | 2 | 0.112 | 4.32 |
| 3 | D9 | 1 | 3.55 | 0.04 |
| 3 | E2 | 2 | 0.464 | 38.58 |
| 3 | F7 | 1 | 1.87 | 0 |
| 3 | G4 | 2* | 0.224 | 10.79 |

### Identification of hermaphrodite genotypes

The hermaphrodite genotypes are those detected as both female and male genotypes. In our dataset, eight genotypes (3.9%) were detected as both female and male. Only three genotypes (1.5% of the total number of female and male genotypes) were supported by significant Psex values and thus considered true hermaphrodites (Table 1 and Fig. 3). The R021 genotype was the most frequent female genotype (see above). It has been also detected in five ascocarps as males and once as an ectomycorrhiza (Fig. 3A). The R060 genotype was found five times as the female genotype, once as a male and once as an ectomycorrhiza (Fig. 3B). Finally, the R068 was detected three times as the female genotype, once as a male, and once as an ectomycorrhiza (Fig. 3C). These three hermaphrodite genotypes were detected as male and/or female in 55 ascocarps (27%). Conversely, 135 genotypes were detected only as male (98%) and 70 genotypes only as female (92%).

## Discussion

In this study, ascocarps, ectomycorrhizas, and soil cores were harvested over a 5-year period under *T. melanosporum* productive trees in a truffle orchard. The female and male genotypes of 238 and 206 ascocarps, respectively, were successfully obtained. The genotypic diversity was higher for male genotypes than female genotypes with numerous small size genotypes suggesting an important annual turnover. However, a few larger and perennial female and male genotypes were detected. A pronounced small-scale genetic structure was identified for both female and male genotypes although IBD analysis indicated that the dispersal capacity was similar for both. Most of the genotypes were detected only as female or male, and only three genotypes (1.5%) were found as being both female and male (hermaphrodite). These data allowed an *in situ* comparison of the temporal dynamic, genetic structure, and diversity of both female and male genotypes providing new information on *T. melanosporum* sexual reproduction.

### Strong small-scale genetic structure for both female and male genotypes

According to the genotypic richness (R) and the Simpson’s diversity index, the genotypic diversity was higher for male than for female genotypes although the gene diversity (1-Qinter index) was similar for both. Large and perennial genotypes were found for both male and female genotypes, but a more rapid annual turnover of male genotypes is suggested by the higher number of small size genotypes (Table 1). A strong IBD and consequently a strong small-scale genetic structure was found for both female and male genotypes with IBD slope value that was not significantly different between sexes (Fig. 4). This result suggests similar dispersal capacities for both female and male genotypes. In fungi, two main dispersal strategies have been well documented: 1) spread of epigeous species' spores by wind and 2) passive dissemination of hypogeous species by animals. The strong IBD observed in our data for both female and male genotypes reflects their low dispersal capacity. This strong genetic structure is characteristic of hypogeous fungi that are expected to have reduced spore dispersal compared to that of epigeous ones (Kretzer et al., 2005). Previous studies already detected a strong small-scale genetic structure for female genotypes (Murat et al., 2013; Taschen et al., 2016), but to our knowledge the male genotype genetic structure has not been investigated to date.

### The strong genetic structure leads to heterozygote deficit

A heterozygote deficit was observed in zygotes (*F*_IS_ value ranging from 0.02 to 0.28), suggesting that female and male genotypes tend to be genetically related in zygotes. Moreover, in nearly 4% of the zygotes, we found female and male genotypes that were homozygous for all microsatellites loci and nearly 50% of the zygotes displayed a kinship coefficient above 0.25 (full-sib level according Loiselle et al. 1995). This strong genetic relatedness within the zygotes is in accordance with the small-scale genetic structure identified and discussed above, suggesting that disseminations over large distances are rare for both female and male genotypes as demonstrated by similar IBD slope values. As already discussed previously, hypogeous fungi rely on animals for spore dissemination, and in most case spores stay in the vicinity of the ascocarps. A more important heterozygote deficit was recently identified in five brulés for *T. melanosporum* with F_IS_ varying from 0.30 to 0.68 and 16.7 to 40% of homozygote zygotes (Taschen et al., 2016). The small-scale genetic structure due to limited dispersal capacities may explain the observed heterozygote deficit. An existing genetic barrier impeding mating of unrelated female and male genotypes is unlikely since 26% of the zygotes have a negative kinship coefficient.

### Germinating aseospores could act as male genotypes

For both female and male genotypes, we observed similar features with both co-occurrence of small size genotypes, often detected transiently as a single ascocarp, and larger perennial genotypes. These results suggest that for both female and male genotypes, there is a mix of new genotypes recruited from ascospores and perennial genotypes that have been disseminated by vegetative propagation. However, male genotypes presented a higher genotypic diversity and were less perennial than female ones, suggesting that ascospore recruitment is more important for male than for female genotypes. As already proposed by Taschen and colleagues (2016), it is therefore tempting to hypothesize that most of the male genotypes originate from germinating ascospores whose mycelium does not survive after sexual reproduction. Indeed, sexual spores have been proposed as male gametes for ascomycetes and basidiomycetes (Nieuwenhuis et al., 2011).

The inoculation of ascospores in truffle orchards in order to improve the production of ascocarps has become a common practice (Olivier et al., 2012), but is totally empirical due to lack of scientific background. This practice is not recent since Ciccarello (1564), Bradley (1726) and Buffon (1749) proposed improving truffle production by inoculating pieces of ascocarps under mature trees. The role of ascospores was not directly demonstrated in our study but, in collaboration with truffle growers' dedicated experiments, have been initiated to confirm their role.

### Soil mycelium could also be a reservoir of male genotypes

In our study, 16 male perennial genotypes were found. One of them, genotype R102, was found in 18 ascocarps (8.7%) over four years (Fig. S5). Among these, only three corresponding to true hermaphrodite genotypes were present on ectomycorrhizas. We have made the assumption that these male genotypes have a poor ability to form associations with the host and that they likely survive as free-living mycelium in the soil or are associated with the roots of non-ectomycorrhizal plants. It has been demonstrated that roots of herbaceous plants can host truffle mycelium, but the nature of this interaction (such as colonization of the rhizosphere, endophytism, and endomycorrhiza as with orchids) is unknown (Gryndler et al., 2014). In our study, we detected the presence of both mating types in 16 out of 20 soil cores. This result demonstrates that close to the ectomycorhizas formed by the female genotype, mycelium of opposite mating type are present as mycelium and consequently the male genotype could survive in the soil through vegetative propagation, saprotrophycally, or in association with the roots of non-hosts.

In many ascomycetes, conidia (asexual spores) serve for vegetative propagation or as a male gamete (Nelson, 1996; Maheshwari, 1999). Urban et al. (2004) described the existence of anamorphous structures producing mitotic conidia in soils where *T. borchii* and *T. oligospermum* ascocarps were present. Healy et al. (2013) suggested that Pezizales mitospores, including *Tuber* mitospores, which failed to form ectomycorrhizas, could act as male gametes. Nevertheless, the question of whether conidia act as male gametes in the *Tuber* species in which they have been found remains unanswered and for the moment, conidia were never yet observed in *T. melanosporum.*

### Sexual reproduction strategy in Tuber melanosporum

In our study, a non-random distribution of female genotypes according to their mating type was observed to form large patches from 5 to 20 m^2^ of different genotypes of the same mating type. This result had already been found for *T. melanosporum* (Rubini et al., 2011a; Murat et al., 2013; Taschen et al., 2016) and *T. aestivum* (Molinier et al., 2016). This aggregation was stable over five years despite genotype turnover (Fig. S3). Interestingly, few ascocarps were harvested in the contact zone of either mating type, suggesting that hermaphroditism, or monoecy, is not widespread in *T. melanosporum* (Fig. 2). Indeed, we found only three genotypes that were detected in 27% of the ascocarps as female or male genotypes, which can be considered as true hermaphrodites (scenario 2c in Fig. 2). It is not surprising to identify hermaphrodite genotypes since for heterothallic ascomycete hermaphroditism is the common rule (Glass and Kuldau, 1992; Nieuwenhuis and Aanen, 2012). However, most genotypes were identified as either female (92%) or male (98%) genotype, suggesting a specialisation in one sex leading to subsequent dioecy. In the population of *T. melanosporum* surveyed in the present study, a mix of a few hermaphrodites genotypes with a majority of female and male genotypes (dioecy) co-occurred suggesting trioecy. Trioecy is known in plants (Joseph and Murthy, 2014; Mirski and Brzosko, 2015) and animals (Weeks et al., 2006; Chaudhuri et al., 2011). In fungi, trioecy was reported for the ascomycete *Trieeromyees* (Benjamin, 1986), and it can exist for *F. fujikuroi* (Leslie, 1995), but it does not seem to be a widespread occurrence. In animals, trioecy can occur when environmental conditions change or when a species colonizes a new habitat, leading to a transition from hermaphroditism to dioecy or *viee versa* (Weeks et al., 2006). Trioecy is therefore a transitory status, and in *Caenorhabditis elegans* it seems evolutionarily unstable (Chaudhuri et al., 2011). In heterothallic ascomycetes, hermaphroditism could be the ancestral status since it is expected for most of the species (Glass and Kuldau, 1992). It is therefore tempting to hypothesize that hermaphroditism has been lost in *T. melanosporum* in order to favour female and male genotypes. But unfortunately, without *in vitro* tests, which are not yet available for *T. melanosporum,* the likelihood of hermaphroditism *versus* dieocy cannot be formally demonstrated.

In conclusion, sexual dimorphism could be more frequent in fungi than expected, and progress in genome sequencing could allow for its investigation. Indeed, in contrast to animals and plants, dioecious fungi often are morphologically similar, and sexual dimorphism can be detected only at genomic or agene regulation levels (Samils et al., 2013). The understanding of economically interesting fungal species' sexual reproduction (such as those producing edible mushrooms) is a major issue for a better control of their life cycles, cultivation, and domestication. It appears that re-inoculation of planted *T. melanosporum* mycorrhizal trees with ascospores leads to an increased number of male gametes (Taschen et al., 2016), improving their dispersal, and favouring the turnover of ectomycorrhizas producing female gametes. Consequently, the re-inoculation of spores is considered as one of the management strategies aiming to decrease the time of appearance for the first fructifications after plantation and increase the number of ascocarps after the initiation of the sexual reproduction.

## Experimental procedures

### Truffle orchard and sampling

Ascocarps and ectomycorrhizas were sampled at a long-term experimental site located at Rollainville in north-eastern France. The work site has been described in a previous work by Murat *et al.* (2013). Samples and trees were identified by a letter and number and mapped on a grid of 1 m × 1 m squares set up with camping pickets (Fig. S1). Three different grids were made to identify three areas (areas 1–3) that cover all of the productive zones of the plantation (Fig. S1).

As described by Murat *et al.* (2013), the sampling started under trees F10, F11 and E10 (area 1) in the 2010-2011 season and under the trees A11, A12, B11 (area 3) and D11 (area 2) in the 2011-2012 season (Fig. S1). The mature truffles were systematically harvested during the production season with help from a well-trained dog, and at the time of harvest, they were precisely mapped on the grid with 5 cm precision. The ascocarps were then washed to remove soil particles and stored at −20°C for molecular analysis.

During this five-year interval, two ectomycorrhizal samplings were done. The first one was done under F10/F11/E10 trees (area 1 in Fig. S1) in spring 2011 (season 1), and results on that sampling were published by Murat et al. (2013). The second one was done in November 2014 (season 5), and 45 root samples were sampled all over the truffle orchard (Fig. S1). Root samples were harvested from the first 10 cm of soil and mapped on the same grid used for the positioning of the ascocarps. The ectomycorrhizas were carefully retrieved from the soil and washed in water under a dissecting microscope. From each root sample, the *T. melanosporum* ectomycorrhizas were identified as described by Zambonelli et al. (1993) and Rauscher et al. (1995), and stored individually in microcentrifuge tubes at −20°C for molecular analyses.

In order to investigate the distribution and abundance of both mating types in the soil, 20 soil cores were harvested in May 2015 (Fig. S1) at a 10-15 cm depth. All plant debris, stones, and roots were discarded from the soil samples, and samples were kept at −20°C for DNA extractions.

### DNA extractions

Genomic DNA from the gleba (inner part of the ascocarps), and ectomycorrhizal tips were extracted by using the DNeasy Plant Mini Kit (Qiagen SA, Courtaboeuf, France), following the manufacturer’s instructions. According to Paolocci and colleagues (2006), when using this protocol, only the haploid female genotype DNA is isolated since spores are not disrupted. To have access to male genotype from each ascocarp, a mixture of spores was isolated, and their DNA was extracted as described below. Thin slices of each ascocarp were put onto a layer of water in a Petri dish to let spores be released into the water in order to isolate the pool of spores. The liquid was collected in a 1.5 mL tube and centrifuged at 14,000 rpm for 5 min.

The supernatant was then discarded in order to obtain a mixture of asci and spores. DNA from the isolated mixture of spores from each ascocarp was extracted as described in Rubini et al. (2011b) with some modifications. First, to each pool of spores, we added 300 mL of NTE buffer (Tris-HCl 200 mM, NaCl 250 mM EDTA 25 mM), and two tungsten beads after which the spores were disrupted with a Tissue Lyser (Qiagen) for 10 minutes at 30 Hz. The tubes were then centrifuged at 14,000 rpm for 10 min, and the recovered supernatant was added to a new tube. Thirty microliters of NaAc (3M) and 330 μL isopropanol were added, and those tubes were mixed and centrifuged again at 14,000 rpm for 10 min. After discarding the supernatant and cleaning the pellet with 200 μL of ethanol (70%), the pellet (DNA) was recovered in 50 μL TE. DNA extracts were stored at −20°C.

Total DNA from soil samples was extracted using the Power Soil^®^ DNA extraction Kit (MoBio, Laboratories, Carlsbad, CA) following manufacturer’s protocol and stored at −20°C for molecular analysis.

### Molecular genotyping

All extracted DNA samples were amplified using species-specific *T. melanosporum* primers (Paolocci *et al.* 1999) in order to check the species and DNA quality.

All of the samples in which *T. melanosporum* identity was confirmed were genotyped by using primer pairs corresponding to ten microsatellite markers (Tm16_ATA12, Tm241_TAA17, Tm2_TAT15, Tm98_TAT15, Tm112_TAT19, Tm9_ATCA12, Tm1_ATTG18, Tm75_GAAA14, Tm22_CCTCAT17 and Tm269_TGTTGC15) as described by Murat et al. (2011; 2013). According to Murat et al. (2011), the use of ten microsatellites allows to reach a genotypic diversity of 0.99. The genotyping was done by the INRA plateform Gentyane (Clermont-Ferrand). The *T. melanosporum* mating types of all the samples were analysed using specific primers for either the *MAT1-1-1* or the *MAT1-2-1* genes using the PCR conditions described by Rubini et al. (2011b). Hereafter, according to Rubini et al. (2011b), the mating types are termed *MAT1-1* and *MAT1-2.*

From the mix of DNA spores, we obtained genotypes of zygote (the mating of female and male gametes for each ascocarp) from which the male genotype was deduced by subtraction of the corresponding female genotype.

The mycelium from soil of both mating types was quantified by quantitative real time PCR (qPCR) using a protocol developed from the international patent (EP2426215 A1) in a confidential research program with ALCINA sarl (Montpellier, France). qPCR reactions were carried out with a StepOne Plus^TM^ Real-Time PCR System machine provided with the StepOne software v. 2.3 (Life Technologies, Carlsbad, CA). Two standard curves (R^2^=0.99; Eff = 96.99% and R^2^=0.99; Eff = 99.67% for *MAT1-1* and *MAT1-2,* respectively) were obtained, as described in Parladé et al. (2013), by mixing a known amount of soil harvested in a cereal field close to the truffle orchard (in which the absence of *T. melanosporum* was confirmed) with a known amount of fresh immature ascocarp of *T. melanosporum* belonging to one or the other mating type. DNA from the mixture was extracted as all the other soil samples and serial tenfold dilutions were done to obtain a standard curve for each mating type. Absolute quantification of mating types in soil samples was obtained by interpolating their threshold cycle (Ct) values on the corresponding standard curve.

### Data analyses

All the samples with null alleles were discarded from our analyses. MLGsim 2.0 (Stenberg et al., 2003) was used for the multilocus genotype identification and the calculation of the likelihood (PSex) that copies of multilocus genotypes result from sexual reproduction or clonal spread. The threshold value (<0.05) for testing the significance of the PSex for each genotype was estimated using 1000 simulations. When the PSex values fell below the threshold value (<0.05), it was concluded that identical genotypes originated from the same genet (clonal multiplication).

In order to analyse clonal and spatial genetic structure of this *T. melanosporum* population, different clonal related indices were estimated. Using the RClone package (Bailleul et al., 2016), the Simpson's diversity index modified for finite sample sizes (1-D) was used as an estimation of genotypic diversity to calculate if two randomly selected samples from the population have the same genotype. The genotypic richness R was also calculated as (G-1)/(N-1) where G is the number of genotypes and N the number of samples (Dorken and Eckert, 2001) The clonal subrange representing the spatial scale over which genetic structure is affected by clonal processes (Alberto et al., 2005) was also calculated. Using Geneclone software (Arnaud-Haond and Belkhir 2007), the aggregation index (Ac) was calculated to test for the existence of spatial aggregation for mating types Arnaud-Haond et al., 2007). Ac index ranges from 0 (when the probability of identity between nearest neighbors does not differ from the average one) to 1 (when all nearest neighbors preferentially share the same genotype). The statistical significance of the Ac index was tested against the null hypothesis of spatially random distribution using a re-sampling approach based on 1000 permutations. The Ac was calculated using culled data for female genotypes and after consideration of the five seasons. In order to study a possible heterozygote deficit, the inbreeding coefficient (*Fis*) was calculated as a difference of observed and expected heterozygosity, using the RClone package. SPAGeDi 1.3 software (Hardy and Vekemans, 2002) was used to estimate the kinship coefficient of Loiselle et al. (1995), as estimator of the genetic relatedness between female and male genotypes in zygotes. Unbiased gene diversity (He=1-Qinter) and isolation by distance (IBD) analysis were obtained with GenePop software v4.2 online (Rousset, 2008). The significance of IBD was tested using Mantel test with 10,000 permutations. All values were calculated for all the data and also separately for female, male, and ECM data. A culled dataset was constructed to reduce the bias due to sampling the same genotype several times. For each season, the isobarycentre was considered for samples sharing the same genotype and having significant Psex. If two samples with the same genotypes were found in different areas of the truffle orchard (areas 1–3), they were separated. Similarly, we did not calculate the isobarycentre with samples sharing the same genotype from different years.

All the maps were obtained with a dedicated python program developed for this study. The mapping program can be downloaded using a Unix terminal and the following command: git clone https://git.igzor.net/inra/iam_mapping.git

## Acknowledgments

The French National Research Agency (ANR) as part of the "Investissements d'Avenir" program (ANR-11-LABX-0002-01, Lab of Excellence ARBRE) and ANR SYSTERRA SYSTRUF (ANR-09-STRA-10) financed this study. ALCINA also contributed by cofinancing the ClimaTruf project. We are grateful to Dr Francesco Paolocci and its team in Perugia (Italy) for providing us the protocol for DNA extraction from a mix of spores and for the constructive discussion. The authors would also like to thank Prof Fabienne Malagnac for the critical reading of the first version of the manuscript. The dog Biella hunted the ascocarps in the truffle orchard. We have no conflict of interest to declare.

## References

Alberto, F., Gouveia, L., Arnaud-Haond, S., Pérez-Lloréns, J. L., Duarte, C. M. and Serrão, E. A. (2005), Within-population spatial genetic structure, neighbourhood size and clonal subrange in the seagrass *Cymodocea nodosa*. Mol Ecol 14: 2669–2681.

Arnaud-Haond, S. and Belkhir, K. (2007) Genclone: a computer program to analyse genotypic data, test for clonality and describe spatial clonal organization. Mol Ecol Notes 7: 15–17.

Arnaud-Haond, S., Duarte, C.M., Alberto, F., and Serrão, E.A. (2007) Standardizing methods to address clonality in population studies. Mol Ecol 16: 5115–5139.

Bailleul, D., Stoeckel, S., and Arnaud-Haond, S. (2016) RClone: a package to identify MultiLocus Clonal Lineages and handle clonal data sets in R. Methods Ecol Evol doi:10.1111/2041-210X.12550.

Benjamin, R.K. (1986) Laboulbeniales on semiaquatic Hemiptera. V. Triceromyces: with a description of monoecious-dioecious dimorphism in the genus. Aliso 11: 245–278.

Bradley, R. (1726) New Improvements of Planting and Gardening: Both Philosophical and Practical: In three parts. London, UK: Printed for W. Mears.

Buffon, G., Sonnini, C.S., and Latreille, P. (1749) Histoire Naturelle Des Plantes. Tome Troisième. Plantes Cryptogames, Des Champignons. In Histoire Naturelle Générale et Particulière Avec La Description Du Cabinet Du Roi (1749 à 1789). Paris, France: Imprimerie Royale.

Chaudhuri, J., Kache, V., and Pires-daSilva, A. (2011) Regulation of Sexual Plasticity in a Nematode that Produces Males, Females, and Hermaphrodites. Curr Biol 21: 15481551.

Chen, J., Murat, C., Oviatt, P., Wang, Y., and Tacon, F.L. (2016) The Black Truffles *Tuber melanosporum* and *Tuber indicum*. In, Zambonelli,A., Iotti,M., and Murat,C. (eds), True Truffle (Tuber spp.) in the World, Soil Biology. Springer International Publishing, pp. 19–32.

Chevalier, G. and Grente, J. (1978) Application pratique de la symbiose ectomycorhizienne: production a grande echelle de plants mycorhizes par la truffe (Tuber melanosporum Vitt.). Mushroom Sci 10: 483–505.

Ciccarello, A. (1564) Opusculum De Tuberibus. Ludouici Bozetti (eds.). Pavia, Italy.

Dorken, M.E. and Eckert, C.G. (2001) Severely reduced sexual reproduction in northern populations of a clonal plant, Decodonverticillatus (Lythraceae). J Ecol 89: 339–350.

Douhan, G.W., Vincenot, L., Gryta, H., and Selosse, M.-A. (2011) Population genetics of ectomycorrhizal fungi: from current knowledge to emerging directions. Fungal Biol 115: 569–597.

Ellstrand, N.C. and Roose, M.L. (1987) Patterns of Genotypic Diversity in Clonal Plant Species. Am J Bot 74: 123–131.

Glass, N.L., and Kuldau, G.A. (1992) Mating Type and Vegetative Incompatibility in Filamentous Ascomycetes. Ann Rev Phytopathol 30: 201–224.

Gryndler, M., Černá, L., Bukovská, P., Hršelová, H., and Jansa, J. (2014) *Tuber aestivum* association with non-host roots. Mycorrhiza 24: 603–610.

Hardy, O.J. and Vekemans, X. (2002) SPAGeDi: a versatile computer program to analyse spatial genetic structure at the individual or population levels. Mol Ecol Notes 2: 618–620.

Healy, R.A., Smith, M.E., Bonito, G.M., Pfister, D.H., Ge, Z.W., Guevara, G.G. et al. (2013) High diversity and widespread occurrence of mitotic spore mats in ectomycorrhizal Pezizales. Mol Ecol 22: 1717–1732.

Joseph, K.S., and Murthy, H.N. (2014) Sexual system of *Qareinia indiea* Choisy: geographic variation in trioecy and sexual dimorphism in floral traits. Plant Syst Evol 301: 1065-1071.

Kretzer, A.M., Dunham, S., Molina, R., and Spatafora, J.W. (2005) Patterns of vegetative growth and gene flow in *Rhizopogon vinicolor* and *R. vesionlosns* (Boletales, Basidiomycota). Mol Ecol 14: 2259–2268.

Le Tacon, F., Rubini, A., Murat, C., Riccioni, C., Robin, C., Belfiori, B. et al. (2015) Certainties and uncertainties about the life cycle of the Perigord black truffle *(Tnber melanosporum* Vittad.). Ann For Sci 73: 105–117.

Leslie, J.F. (1995) *Gibberella fujikuroi:* available populations and variable traits. Can J Bot 73: 282–291.

Leslie, J.F., and Klein, K.K. (1996) Female Fertility and Mating Type Effects on Effective Population Size and Evolution in Filamentous Fungi. Genetics 144: 557–567.

Loiselle, B.A., Sork, V.L., Nason, J., and Graham, C. (1995) Spatial genetic structure of a tropical understory shrub, Psychotria officinalis (Rubiaceae). Am J Bot 1420–1425.

Maheshwari, R. (1999) Microconidia of *Nenrospora erassa*. Fnngal Genet Biol 26: 1–18.

Martin, F., Kohler, A., Murat, C., Balestrini, R., Coutinho, P.M., Jaillon, O. et al. (2010) Périgord black truffle genome uncovers evolutionary origins and mechanisms of symbiosis. Nature 464: 1033–1038.

Mirski, P., and Brzosko, E. (2015) Are hermaphrodites better adapted to the colonization process in trioecious populations of *Salix myrsinifolia*? Acta Soc Bot Polo 84: 167–175.

Molinier, V., Murat, C., Baltensweiler, A., Büntgen, U., Martin, F., Meier, B. et al. (2016) Fine-scale genetic structure of natural *Tnber aestivnm* sites in southern Germany. Myeorrhiza 26: 85–907.

Murat, C. (2015) Forty years of inoculating seedlings with truffle fungi: past and future perspectives. Myeorrhiza 25: 77–81.

Murat, C., Riccioni, C., Belfiori, B., Cichocki, N., Labbé, J., Morin, E. et al. (2011) Distribution and localization of microsatellites in the Perigord black truffle genome and identification of new molecular markers. Fungal Genet Biol 48: 592–601.

Murat, C., Rubini, A., Riccioni, C., De la Varga, H., Akroume, E., Belfiori, B. et al. (2013) Fine-scale spatial genetic structure of the black truffle (*Tuber melanosporum*) investigated with neutral microsatellites and functional mating type genes. New Phytol 199: 176–187.

Nelson, M.A. (1996) Mating systems in ascomycetes: a romp in the sac. Trends Genet 12: 69–74.

Nieuwenhuis, B.P.S., and Aanen, D.K. (2012) Sexual selection in fungi. J Evol Biol 25: 2397–2411.

Nieuwenhuis, B.P.S., Debets, A.J.M., and Aanen, D.K. (2011) Sexual selection in mushroomforming basidiomycetes. Proe R Soc Lond B Biol Sci 278: 152–157.

Olivier, J., Savignac, J., and Sourzat, P. (2012) Truffe et trufficulture. Périgueux, France: FANLAC Editions.

Paolocci, F., Rubini, A., Granetti, B., and Arcioni, S. (1999) Rapid molecular approach for a reliable identification of *Tuber* spp. ectomycorrhizae. FEMS Microbiol Ecol 28: 23–30.

Paolocci, F., Rubini, A., Riccioni, C., and Arcioni, S. (2006) Reevaluation of the Life Cycle of *Tuber magnatum*. Appl Environ Microbiol 72: 2390–2393.

Parladé, J., De la Varga, H., De Miguel, A., Sáez, R., and Pera, J. (2013) Quantification of extraradical mycelium of *Tuber melanosporum* in soils from truffle orchards in northern Spain. Myeorrhiza 23: 99–103.

Rauscher, T., Agerer, R., and Chevalier, G. (1995) Ektomykorrhizen von *Tuber melanosporum, Tuber mesentericum und Tuber rufum* (Tuberales) *an Corylus avellana*. Nova Hedwigia 61: 281–322.

Rousset, F. (2008) GENEPOP’007: a complete re-implementation of the genepop software for Windows and Linux. Mol Ecol Resour 8: 103–106.

Rubini, A., Belfiori, B., Riccioni, C., Arcioni, S., Martin, F., and Paolocci, F. (2011a) *Tuber melanosporum:* mating type distribution in a natural plantation and dynamics of strains of different mating types on the roots of nursery-inoculated host plants. New Phytol 189: 723–735.

Rubini, A., Belfiori, B., Riccioni, C., Tisserant, E., Arcioni, S., Martin, F., and Paolocci, F. (2011b) Isolation and characterization of MAT genes in the symbiotic ascomycete *Tuber melanosporum*. New Phytol 189: 710–722.

Samils, N., Gioti, A., Karlsson, M., Sun, Y., Kasuga, T., Bastiaans, E. et al. (2013) Sex-linked transcriptional divergence in the hermaphrodite fungus *Neurospora tetrasperma*. P Roy Soc Lond B Biol Sci 280: 20130862.

Stenberg, P., Lundmark, M., and Saura, A. (2003) MLGsim: a program for detecting clones using a simulation approach. Mol Ecol Notes 3: 329–331.

Taschen, E., Rousset, F., Sauve, M., Benoit, L., Dubois, M.-P F. Richard, F. and Selosse, MA. (2016). How the truffle got its mate: insights from genetic structure in spontaneous and planted Mediterranean populations of *Tuber melanosporum*. Mol Ecol doi:10.1111/mec.13864.

Urban, A., Neuner-Plattner, I., Krisai-Greilhuber, I., and Haselwandter, K. (2004) Molecular studies on terricolous microfungi reveal novel anamorphs of two Tuber species. Mycol Res 108: 749–758.

Vekemans, X. and Hardy, O.J. (2004) New insights from fine-scale spatial genetic structure analyses in plant populations. Mol Ecol 13: 921–935.

Weeks, S.C., Sanderson, T.F., Reed, S.K., Zofkova, M., Knott, B., Balaraman, U. et al. (2006) Ancient androdioecy in the freshwater crustacean Eulimnadia. P Roy SOC LondB Biol Sci 273: 725–734.

Zambonelli, A., Salomoni, S., and Pisi, A. (1993) Caratterizzazione anatomo-morfologica delle micorrize di *Tuber* spp. su *Qnercnspnbescens* Will. Micol Ital 3: 73–90.

